# PhotoFiTT: A Quantitative Framework for Assessing Phototoxicity in Live-Cell Microscopy Experiments

**DOI:** 10.1101/2024.07.16.603046

**Authors:** Mario Del Rosario, Estibaliz Gómez-de-Mariscal, Leonor Morgado, Raquel Portela, Guillaume Jacquemet, Pedro M. Pereira, Ricardo Henriques

**Affiliations:** Optical cell biology group, Instituto Gulbenkian de Ciência, Oeiras, Portugal; Abbelight, Cachan, France; Intracellular Microbial Infection Biology Laboratory, Instituto de Tecnologia Química e Biológica António Xavier, Universidade Nova de Lisboa, Portugal; Turku Bioimaging, University of Turku and Åbo Akademi University, Turku, Finland; Faculty of Science and Engineering, Cell Biology, Åbo Akademi University, Turku, Finland; InFLAMES Research Flagship Center, Åbo Akademi University, Turku, Finland; Turku Bioscience Centre, University of Turku and Åbo Akademi University, 20520, Turku, Finland; MRC Laboratory for Molecular Cell Biology, University College London, London, United Kingdom

**Author notes:** Correspondence: (P. M. Pereira), (R. Henriques). Equally contributed authors (Mario Del Rosario, Estibaliz Gómez-de-Mariscal).

**Keywords:** phototoxicity, microscopy, machine learning, cell division, cellular activity

## Abstract

Phototoxicity in live-cell fluorescence microscopy can compromise experimental outcomes, yet quantitative methods to assess its impact remain limited. Here we present PhotoFiTT (Phototoxicity Fitness Time Trial), an integrated framework combining a standardised experimental protocol with advanced image analysis to quantify light-induced cellular stress in label-free settings. PhotoFiTT leverages machine learning and cell cycle dynamics to analyse mitotic timing, cell size changes, and overall cellular activity in response to controlled light exposure. Using adherent mammalian cells, we demonstrate PhotoFiTT’s ability to detect wavelength- and dose-dependent effects, showcasing that near-UV light induces significant mitotic delays at doses as low as 0.6 *J/cm*^2^, while longer wavelengths require higher doses for comparable effects. PhotoFiTT enables researchers to establish quantitative benchmarks for acceptable levels of photodamage, facilitating the optimisation of imaging protocols that balance image quality with sample health.

## Main

Live-cell fluorescence microscopy has revolutionised our ability to study dynamic biological processes in near-native conditions (1). However, the intense illumination required can induce phototoxicity, disrupting cellular functions (2– 5). These effects, often subtle and cumulative, can lead to significant changes in cell behaviour, organelle integrity, and developmental processes (6–9). Yet, establishing optimal imaging protocols that balance high-quality data acquisition with minimal biological interference remains a complex challenge (10). Traditional methods for assessing phototoxicity include viability assays and morphological observations. Historically, photobleaching has been used as a proxy reporter of phototoxicity, but with the advent of highly photostable fluorescent proteins like mStayGold (11), this approach has become less reliable. Importantly, tolerance to photodamage varies across specimens and is influenced by damage severity, complicating the development of replicable imaging protocols (12, 13). Previous studies have offered valuable quantitative assessments of phototoxicity (3, 10, 14), but a critical need remains for standardised and generalisable methods able to quantitatively link cell damage to high-intensity light exposure across different fluorescence microscopy techniques. Addressing this gap, we introduce PhotoFiTT (Phototoxicity Fitness Time Trial), a quantitative imaging-based framework designed to assess phototoxicity effects on cellular behaviour in live-cell microscopy experiments (**Figure 1**). PhotoFiTT provides a rigorous, labelfree, and quantitative approach to evaluate phototoxicity, enhancing the reliability and reproducibility of live-cell imaging studies. It is designed as an integrated framework comprising a standardised experimental protocol and an advanced image analysis pipeline. PhotoFiTT leverages the predictable nature of cell division as a “biological clock” to quantify phototoxicity-induced perturbations (**Figure 1**). The framework uses low-illumination brightfield microscopy to monitor cell populations that have been exposed to controlled light damage events. It employs deep learning algorithms, including virtual staining and cell segmentation, to aid in tracking abnormal light-induced cellular behaviour. PhotoFiTT employs three key measurements: 1) **Mitosis monitoring:** assess delays in cell division by mapping mitotic cell rounding in time (**Figure 1a**); 2) **Cell size dynamics:** track changes in cell size to discriminate between cell division and cell cycle arrest (**Figure 1b**); 3) **Cellular activity:** quantify subcellular changes over time as a measure of overall cellular health (**Figure 1c**). By analysing these factors, PhotoFiTT enables researchers to extract numerical constraints for optimising live-cell-compatible illumination conditions.

**Fig. 1.**
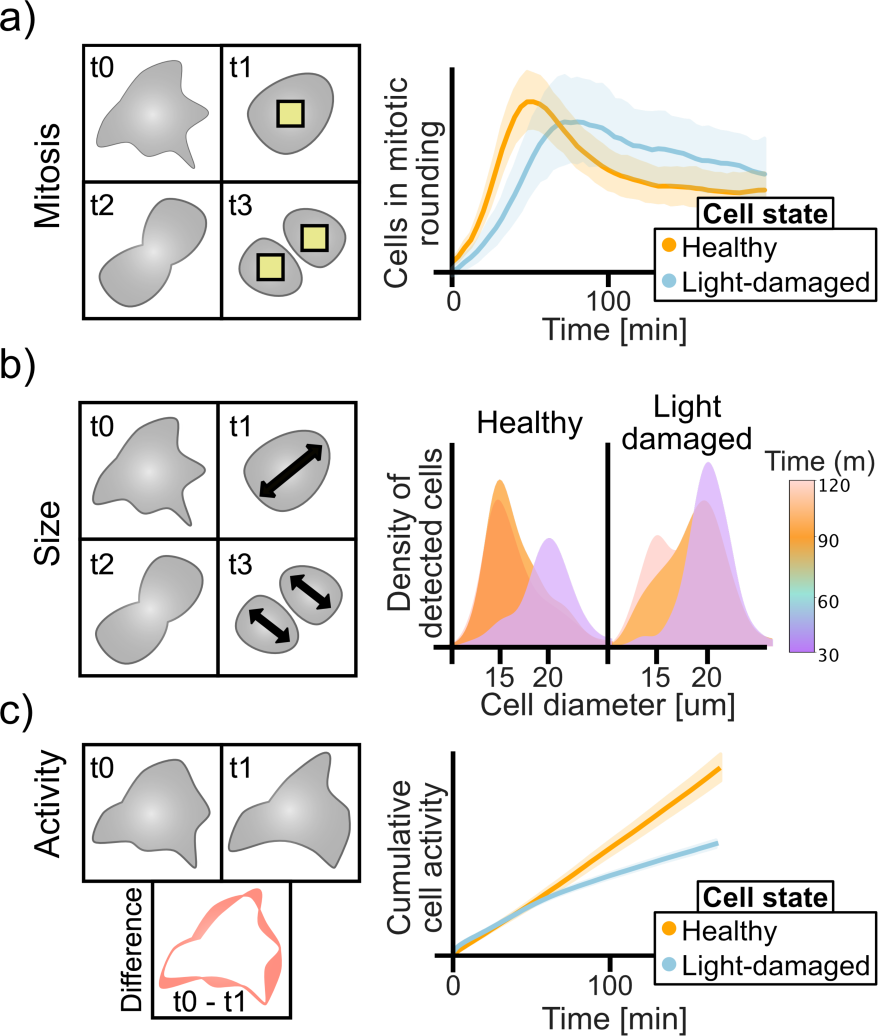
Phototoxicity fitness time trial (PhotoFiTT) assay. PhotoFiTT integrates a biological imaging assay with an image analysis workflow, using cellular processes and mitotic cycles as natural timers to track consistent patterns of cell behaviour. It detects deviations in these patterns caused by light exposure, enabling the quantification of phototoxic effects in fluorescence microscopy. The workflow analyses three cellular features: a) Mitosis: Identifies mitotic rounding (yellow squares), typically 30-50 minutes post-G2 exit. Phototoxicity alters rounding time distributions (orange: normal, blue: photodamaged). b) Size: Tracks cell diameters (black arrowheads) through division. Mother cells (purple) transition to daughter cells (orange) approximately 60 minutes post G2 exit. Phototoxicity delays this transition (light orange). c) Activity: Quantifies sub-cellular changes between frames (red outlines). Normal cells show increased post-division activity (orange, around 60 minutes), while photodamaged cells (blue) exhibit reduced activity.

To implement PhotoFiTT (**Figure 2a**), researchers can follow these key steps: 1) Synchronise cells using the CDK-1 inhibitor RO-3306, which arrests cells in the G2/M interface, or study unsynchronised cells that can be tracked from mitotic rounding until division; 2) Expose cells to a light irradiation event replicating the illumination pattern of the imaging experiment to be analysed for phototoxity; 3) Identify cells undergoing mitotic rounding and division (**Figure 2b**) using a Stardist (15) deep learning model pursposely-trained for brightfield round cell detection; 4) Analyse three key parameters: mitotic timing, cell diameter dynamics, and cellular activity. **Video S1-2** provide in-detail tutorials to reproduce these procedures. We conducted a comprehensive analysis of phototoxicity using Chinese Hamster Ovary (CHO) cells as a model system. CHO cells have the benefit of having a short doubling time (14-17 hours), allowing for efficient experimental timelines, while being a highly popular cell line for microscopy and cell research. Both synchronised and unsynchronised cell populations were exposed to varying doses of near-UV (385 *nm*), blue (475 *nm*), and red (630 *nm*) light to assess wavelength-dependent effects. In synchronised cells, near-UV irradiation induced significant dose-dependent delays in mitotic timing, with effects detectable at doses as low as 0.6 *J/cm*^2^ (**Figure 2c-d, Video S3**). Notably, at 60 *J/cm*^2^, the characteristic peak in mitotic rounding became almost imperceptible, indicating widespread cell cycle arrest. To validate that these effects were primarily due to light exposure rather than the potential light sensitivity of the CDK1 inhibitor used for synchronisation, we conducted parallel experiments with unsynchronised cells. In unsynchronised cell populations exposed to near-UV light, we observed a dose-dependent decrease in dividing cells and a simultaneous increase in arrested cells (**Figure S1a**). Consistent with our findings in synchronised populations, the time required for cells to divide after entering mitotic rounding increased with higher light doses (**Figure S1b**). These results confirm that the observed effects were primarily attributable to light exposure rather than light sensitivity increase by the synchronisation drug. Notably, synchronised populations exhibited higher sensitivity to near-UV-induced damage compared to their unsynchronised counterparts. This increased susceptibility was evidenced by a higher proportion of apoptotic cells, as identified by a SYTOX-based cell viability assay (**Figure S1c**). This observation can be attributed to the fact that synchronised cells were uniformly arrested at the G2/M checkpoint, a critical stage immediately following DNA replication and preceding mitosis. Near-UV radiation is known to induce DNA damage (12), and cells in G2 phase are particularly vulnerable due to the presence of fully replicated chromosomes and the imminent onset of mitosis. The accumulation of DNA damage at this stage can trigger cell cycle arrest or apoptosis more readily than in unsynchronised populations, where cells are distributed across various cell cycle phases (16). While near-UV irradiation demonstrated the most pronounced effects, longer wavelengths also induced cellular stress at higher doses. Cells exposed to 475 *nm* and 630 *nm* light began showing comparable delays to those observed with near-UV light (i.e., mitotic rounding time points above 60 *min*) only at the highest tested dose of 60*J/cm*^2^ (**Figure 2d**). This finding highlights a clear wavelength dependence in phototoxicity, with longer wavelengths requiring substantially higher doses to induce comparable photodamage. PhotoFiTT’s analysis of cell size dynamics revealed that near-UV exposure delayed the appearance of daughter cells in a dose-dependent manner (**Figure 2e**). Under control conditions, mother cells (*≥* 20 *µ*m diameter) divide into daughter cells (*≥* 15 *µ*m diameter) within 50 *≠* 75 minutes post-synchronisation. Near-UV light exposure induced dosedependent delays in this process, with high doses (60*J/cm*^2^) delaying daughter cell appearance by up to 120 minutes after synchronisation. Such delay in division and the persistence of mother cells upon time indicates the increase in the number of cells arrested due to photodamage and explains the lost peak in the distribution of mitotic rounding with high doses (60*J/cm*^2^) in **Figure 2c**. Classifying mother and daughter cells according to their size lets PhotoFiTT evaluate cell division delays (**Figure 2f**). Upon challenged with longer wavelengths (475, and 630*nm*) and in agreement with our previous observations (**Figure 2d**), cell division delays are also wavelength-dependant, requiring higher light doses to induce noticeable effects (**Figure 2f**). Beyond acute effects on cell division, PhotoFiTT can also quantify post-mitotic cellular activity, providing insight into long-term impact on cellular behaviour. We observe that higher light doses and shorter wavelengths led to decreased cumulative cellular activity over a 7-hour period (**Figure 2g**). While each of these metrics has its own benefits, tracking the 1-to-2 transition of a mother cell to two resolvable daughter cells provides a highly efficient and straightforward approach to assessing photo-damage effects. This transition can be easily monitored by classifying cell rounding events preand post-division within a short time window (15 *≠* 30 *min*). In control, synchronised populations, approximately 50% of cells transition from resolvable mother to daughter cells within the first 50 minutes post-synchronisation. At a light dose of 6 *J/cm*^2^, the proportion of cells completing the mother-to-daughter transition within this time-frame is reduced to around 40% for 630 *nm* illumination, 30% for 475 *nm*, and only 10% for 405 *nm* (**Figure 2e-f**). Taken together, our results demonstrate the nuanced and continuous nature of phototoxicity. They reveal a spectrum of cellular responses that vary with both light dose and wavelength, underscoring the importance of considering them in experimental design.

**Fig. 2.**
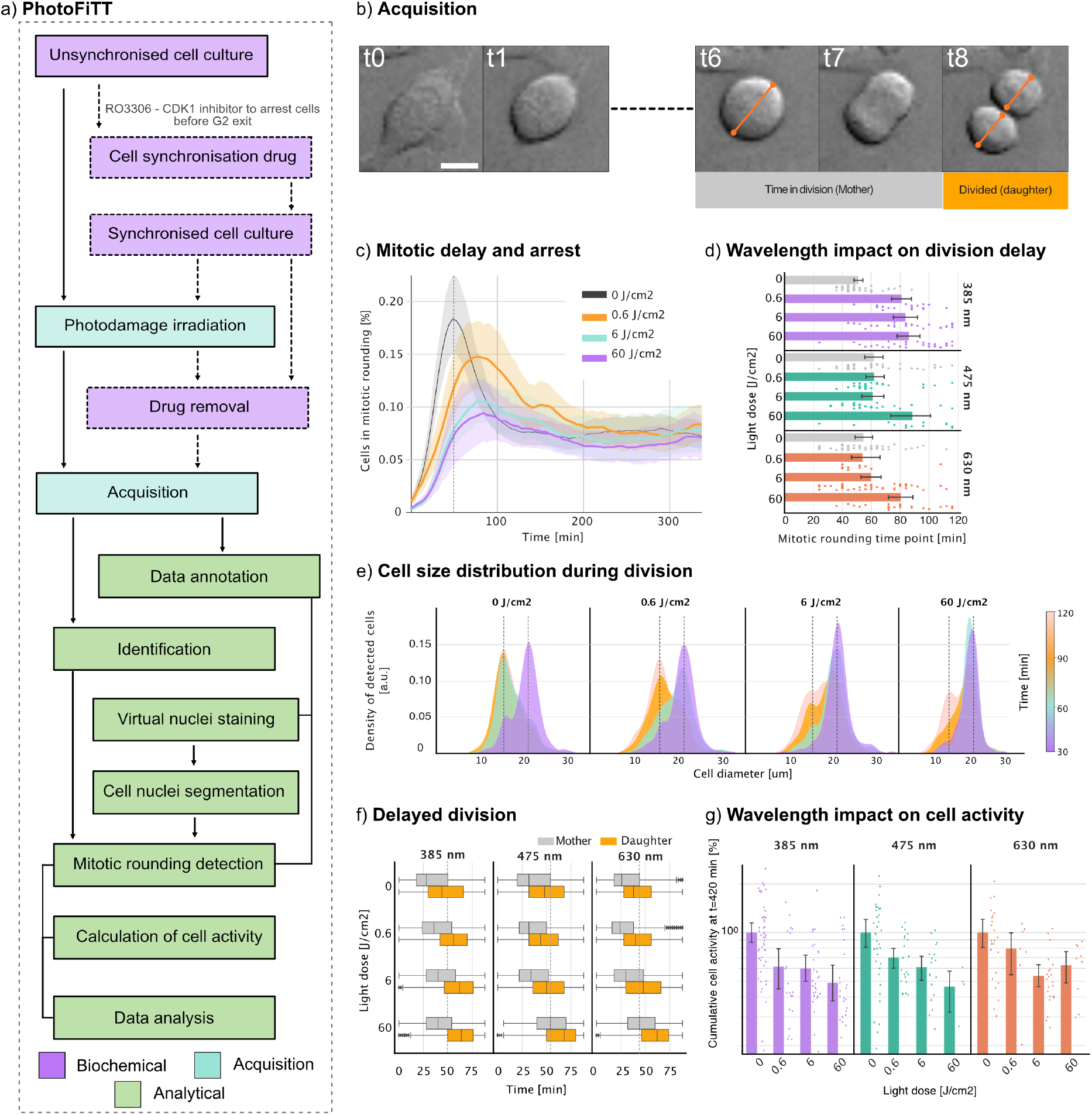
Quantitative phototoxicity assessment with PhotoFiTT. a) PhotoFiTT workflow combines biochemical experiments, microscopy image acquisition and image analysis. Cell cycle synchronisation is induced by a CDK1 inhibitor (RO-3306) arresting cells at the G2/M interface. Photodamage irradiation represents the desired illumination patterns to be studied. Synchronised cells are washed from the drug used prior to acquisition. Time-lapse acquisition starts, using transmitted light, lasting 6 hours. A small subset of the videos is manually annotated to train a cell mitotic rounding identification model. The main experimental time-lapse videos are processed to identify the proportional mitotic rounding with the trained models and calculate the cell activity across time. These measurements are used for the final data analysis. Detailed workflow description **Note S1** and reproducibility guidelines **Note S2** are given in the Supplementary Information and Methods section. b) Time-lapse brightfield imaging of an adherent mammalian cell (Chinese hamster ovary CHO) undergoing mitosis. The mother cell’s diameter approximately doubles that of the resulting daughter cells post-division. Scale bar 20 *µm*. c) Temporal distribution of mitotic cell rounding in synchronised populations exposed to varying doses of 385*nm* (near-UV) light (0.6, 6, and 60 *J/cm*^2^). Exposed populations exhibit a dose-dependent delay in mitotic rounding, manifested as rightward shifts in the distribution peaks. d) Quantification of mitotic rounding delays across different light wavelengths and doses. The control population peaks at *t* 50 minutes, representing the average mitotic rounding time (indicated with a line in b). Exposure to 385*nm* light induces a dose-dependent delay, while 475 *nm* and 630 *nm* wavelengths only cause significant delays at high doses (60*J/cm*^2^). Cell volume nearly halves as mother cells divide into two daughter cells. Low-dose exposure (0.6*J/cm*^2^) delays this transition, while high doses result in heterogeneous populations at later time points (120*min*), indicating asynchronous divisions and potential cell cycle arrest. f) Temporal analysis of mother and daughter cell populations across different wavelengths and doses. All wavelengths induce delays in daughter cell appearance, with 385*nm* light and high doses of 475*nm* and 630*nm* light causing the most pronounced effects. g) Cumulative cell activity over 7*h* hours post-exposure normalised for each replica. All light exposures reduce overall cell activity, with 385*nm* and 475*nm* showing more dose-dependent effects than 630*nm* light. These results collectively demonstrate the wavelength- and dose-dependent impacts of light exposure on cell division dynamics and overall cellular activity.

Our findings reveal that phototoxicity not only induces delays in cell division but also compromises subsequent cellular functions (**Figure S2c-g**), corroborating and extending previous observations. The damage inflicted by shorter wavelengths is more pronounced, likely due to their known direct interaction with DNA (17). Longer wavelengths, while generally less harmful, have been described to exert significant effects on cellular physiology, including alterations in mitochondrial membrane potential, cytoskeletal dynamics, and reactive oxygen species levels (3, 18). The observed physiological responses likely stem from a complex interplay of multiple cellular perturbations, rather than isolated phototoxic effects. This multifaceted nature of phototoxicity underscores the importance of comprehensive assessment approaches in live-cell imaging experiments. A key strength of PhotoFiTT is its ability to reveal the non-binary nature of phototoxicity, demonstrating a continuum of effects that vary with light dose and wavelength in a label free manner. This offers advantages over simple viability assays (**Figure S1c**) and enables fine assessment of specific illumination conditions with minimal interference from the method itself. By iteratively applying PhotoFiTT with different illumination parameters, researchers can identify optimal settings that balance image quality with minimal phototoxicity. This approach allows for fine-tuning of imaging protocols based on quantitative data rather than qualitative assessments. Researchers can systematically vary labels, light doses, wavelengths, and exposure times, then use PhotoFiTT to measure the resulting effects on cell division timing, cell size dynamics, and overall cellular activity. Using PhotoFiTT results to establish quantitative benchmarks for acceptable levels of photodamage, researchers can make informed decisions about imaging parameters and more accurately interpret their results. This is particularly important given the growing recognition of the “reproducibility crisis” in biomedical research (19). Additionally, PhotoFiTT measurements enable comprehensive tracking of cells’ physiology deviations, making it suitable as an early-warning system for cellular stress. It is important to acknowledge limitations of PhotoFiTT. While highly sensitive to major cellular perturbations, it may not detect subtle light-induced events. Its primary strength lies in identifying conditions that significantly impact cell division and overall cellular health without requiring fluorescent labels. PhotoFiTT is designed for adherent, dividing cell lines, which may limit its applicability to other experimental systems. Our results underscore the importance of considering the potentially damaging effects of activation light in super-resolution techniques such as Single Molecule Localisation Microscopy (SMLM) and Single Particle tracking (SPT), which often use illumination in the range of *kW/cm*^2^ for several minutes. For imaging domains where illuminations are well above the 60 *J/cm*^2^ threshold studied here, researchers should carefully consider and mitigate photodamage. Future work could explore the development of accelerated protocols or the integration of PhotoFiTT principles into real-time analysis pipelines to apply corrective measures to avoid cell death. The principles underlying PhotoFiTT could also be extended to other imaging modalities or adapted for specific biological questions, potentially allowing for real-time adjustment of imaging parameters to minimise photodamage during long-term experiments.

## DATA AVAILABILITY

The data obtained in this study, as well as annotated images are available through the BioArchive at https://www.ebi.ac.uk/biostudies/bioimages/studies/S-BIAD1269 and example images to test the code are available through https://zenodo.org/records/12733476 both under CC BY 4.0 license.

## CODE AVAILABILITY

The PhotFiTT computational framework is available at https://github.com/HenriquesLab/PhotoFiTT. We used ZeroCostDL4Mic to train and assess the StarDist and Pix2Pix models: https://github.com/HenriquesLab/ZeroCostDL4Mic. All source code is under an MIT License.

## AUTHOR CONTRIBUTIONS

R.H. and P.M.P. conceived the project. M.DR., E.G.M. P.M.P. and R.H. designed the experimental and analytical pipelines. M.DR. and L.M. performed the live imaging experiments. M.DR., E.G.M. and R.P. acquired the brightfield-fluorescent paired data. M.DR., E.G.M. and L.M. annotated and curated the data. E.G.M. developed the analytical pipeline. M.DR. wrote the Fiji macros. E.G.M and M.DR. analysed the data with inputs from G.J., P.M.P. and R.H.. M.DR. and E.G.M. prepared and tested the GitHub repository and wrote the paper with input from G.J., P.M.P. and R.H. All authors reviewed and refined the manuscript.

## ACKNOWLEDGEMENTS

We thank Kalina Tosheva, Buzz Baum and Caren Norden for discussions regarding experimental workflows. M.DR., E.G.M. and R.H. acknowledge the support of the Gulbenkian Foundation (Fundação Calouste Gulbenkian), the European Research Council (ERC) under the European Union’s Horizon 2020 research and innovation programme (grant agreement No. 101001332) (to R.H.) and funding from the European Union through the Horizon Europe program (AI4LIFE project with grant agreement 101057970-AI4LIFE and RT-SuperES project with grant agreement 101099654-RTSuperES to R.H.). Funded by the European Union. Views and opinions expressed are, however, those of the authors only and do not necessarily reflect those of the European Union. Neither the European Union nor the granting authority can be held responsible for them. This work was also supported by a European Molecular Biology Organization (EMBO) installation grant (EMBO-2020-IG-4734 to R.H.), an EMBO postdoctoral fellowship (EMBO ALTF 174-2022 to E.G.M.), a Chan Zuckerberg Initiative Visual Proteomics Grant (vpi-0000000044 with https://doi.org/10.37921/743590vtudfp to R.H.) and a Chan Zuckerberg Initiative Essential Open Source Software for Science (EOSS6-0000000260). A joint-collaboration between Abbelight and the Instituto Gulbenkian de Ciência kindly supports L.M. R.H. and P.M.P acknowledge funding by Fundação para a Ciência e Tecnologia (FCT, Portugal) through national funds to the Associate Laboratory LS4FUTURE (LA/P/0087/2020, DOI 10.54499/LA/P/0087/2020). P.M.P. acknowledges support from Fundação para a Ciência e Tecnologia (Portugal) project grant (PTDC/BIA-MIC/2422/2020), the R&D unit Mostmicro (UIDB/04612/2020, UIDP/04612/2020), La Caixa Junior Leader Fellowship (LCF/BQ/PI20/11760012) financed by “la Caixa” Foundation (ID 100010434) and by European Union’s Horizon 2020 research and innovation programme under the Marie Skłodowska-Curie grant agreement No 847648, and a Maratona da Saúde award. This study was also supported by the Research Council of Finland (338537 to G.J.), the Sigrid Juselius Foundation (to G.J.), the Cancer Society of Finland (Syöpäjärjestöt; to G.J.), and the Solutions for Health strategic funding to Åbo Akademi University (to G.J.). This research was supported by the InFLAMES Flagship Programme of the Academy of Finland (decision numbers: 337530, 337531, 357910, and 357911).

## EXTENDED AUTHOR INFORMATION

M. Del Rosario: [mariod](https://twitter.com/_mariod)

E. Gómez-de-Mariscal: gomez_mariscal

L. Morgado: [ALeonorMorgado](https://twitter.com/ALeonorMorgado)

R. Portela:

G. Jacquemet: [guijacquemet](https://twitter.com/guijacquemet)

P. M. Pereira: P_Matos_Pereira

R. Henriques: [HenriquesLab](https://twitter.com/HenriquesLab)

## Methods

### Cell lines

CHO (ATCC CCL-61) cells were cultured in DMEM (Gibco) supplemented with either 42 *µ*M gentamicin (Gibco) or 1% penicillin/streptomycin (Gibco) and 10% foetal bovine serum (FBS; Gibco). All cells were grown at 37 ºC in a 5% CO_2_ humidified incubator.

### Cell cultures, synchronisation

Cells were seeded on an 8 well-chambered cover glass (Cellvis) with a total of 6*x*10^4^ cells per well. If cell synchronisation was required, cells were incubated with 10 *nM* RO-3306 (Sigma-Aldrich) to inhibit CDK1 activity for 16 *≠* 18 hours. Unsynchronised cells were seeded using the same protocol and allowed to grow for the same duration.

### Cell fixation and nuclei staining to generate fluorescent and brightfield paired images

Cells grown on a 8 well-chambered cover glass (Cellvis) until 60% confluency were fixed with a 4% paraformaldehyde solution (Electron Microscopy Sciences) for 10 minutes and washed three times with PBS. Subsequently, the samples were incubated with a 0.1*µg/ml* Hoechst 33342 solution for 10 minutes in the dark.

#### A. Excitation light irradiation

Cells were then exposed to excitation light at the Axio Observer 7 (Zeiss) microscope equipped with a Colibri 7 LED illumination (Zeiss). 10 fields of view (FOV) were randomly selected per exposure condition. The exposure time was set for different fields of view in the same well according to the experimental conditions (100 ms corresponding to 0.6*J/cm*^2^, 1 s to 6*J/cm*^2^ and 10 s to 60*J/cm*^2^). The power used for each wavelength line was the following: 84.9 *±* 2.8 *mW* for the 385 *nm* line, 85.2 *±* 0.8 for the 475 *nm*, and 70.6 *±* 0.4 for 630 *nm* (mean *±* standard deviation). Exposed areas were carefully selected so as not to overlap. Immediately after light exposure and before imaging, cells were washed two times with PBS and RO3306 containing media was replaced by phenol-red free Fluorobrite DMEM (Gibco) supplemented with 2 mM GlutaMAX (Gibco), 42 *µ*M gentamicin (Gibco) and 10% FBS (Gibco). The media exchange was done with a custom-made 4-syringe holder designed in TinkerCAD (https://www.tinkercad.com/) and 3D printed using a Prusa MK4 3D printer (.stl file in supplementary material).

#### B. Live-cell imaging and generation of fluorescent and brightfield paired images for training

Live-cell imaging was performed on the Axio Observer 7 (Zeiss) microscope equipped with a Colibri 7 LED illumination (Zeiss), a 20x/0.8 Plan Apochromat objective (Zeiss) and a Prime 95B sCMOS camera (Teledyne Photometrics). Images were acquired every 4 minutes for 8 hours and have a pixel size of 0.55*µm ×* 0.55*µm*. The temperature and CO_2_ levels were maintained at 37 ºC and 5% respectively throughout the excitation light exposure and imaging periods. For fixed samples, cells were imaged using the same objective in a multichannel acquisition mode. Brightfield images were captured alongside fluorescent images using the 385 nm filter to obtain paired brightfield and Hoechst-stained images.

#### C. Live-cell viability

Cell viability was performed by adding 0.1*µM* SYTOX Orange (Thermo) to previously seeded cells and taken to the microscope for live-cell imaging for 12 hours. Live-cell imaging was performed on the Nikon Eclipse Ti2 microscope equipped with a CoolLED pe800 illumination (Nikon), a 20x/0.8 Plan Apochromat objective (Nikon) and an Orca-fusionBT camera (Hamamatsu). 10 fields of view (FOV) were randomly selected per exposure condition and the images were acquired every 4 minutes using the brightfield and TRITC channel in a multichannel acquisition. The temperature and CO_2_ levels were maintained at 37 ºC and 5 %.

#### D. General image processing workflow

As depicted in Figure 2, the image analysis to extract the measurements of interest is composed of: (1) detecting the number of cells in the field of view; (2) detecting the cells in mitotic rounding; (3) estimating the general changes in cell activity. These three steps enable extracting measurements for the numerical quantification of photodamage, such as cell size distribution across time.

##### D.1. Estimation of the number of cells in the field of view using virtual cell nuclei staining and segmentation

The total number of cells in the field of view was computed by first, inferring the cell nuclei from brightfield images using an in-house trained Pix2Pix model (20), and then, segmenting each nucleus with a pre-trained StarDist model (*StarDist-versatile*) provided by the developers as “2D versatile fluo” model and trained on a subset from the Data Science Bowl Kaggle Challenge (21). Using ZeroCostDL4Mic (22), Pix2Pix was trained with paired images of fixed cells stained with Hoechst and imaged with brightfield and widefield fluorescence microscopy. The imaged data was divided into 400 and 132 fields of view for training and testing, respectively. All the input images were preprocessed as follows: a bleach correction to remove the illumination artefacts was computed by applying a large low-pass filter (a wide Gaussian filter) and subtracting it from the original image; then, the image intensity values were normalised with the min-max projection to the [0, 1] range. Both pre-processed input and output images were then normalised using a percentile normalisation of 1% and 99.9% respectively (denoted as Contrast enhancement in the ZeroCostDL4Mic Pix2Pix notebook). During Pix2Pix training, images are also reshaped to have a size of 1024 *×* 1024 pixels, so the images were reshaped to have a pixel size of 0.644 *µu ×* 0.644 *µm*. Pix2Pix was trained from scratch with a patch size of 512 *×* 512 pixels, a batch size of 5 and for 2000 epochs with a learning rate of 0.001 and 1000 more epochs with a linear learning rate decay. The accuracy results for the test dataset are as follows: Structural Similarity Index Measure (SSIM) (0.86), Learned Perceptual Image Patch Similarity (LPIPS) (0.10), Peak Signal-to-Noise Ratio (PSNR) (25.17), Normalised Root Mean Squared Error (NRMSE) (0.13).The output PDF reports of ZeroCostDL4Mic for both the model training and the quality control check are attached to the supplementary material. The Pix2Pix model instance corresponding to the last epoch was chosen to process the first frame of each cell synchronisation video. The output of Pix2Pix was then segmented using the pre-trained StarDist model. The total number of individually segmented nuclei was used as the number of cells in the field of view with an average of 126 cells per FOV.

##### D.2. Cell mitosis detection with StarDist

A 2D StarDist (15) model -*StarDist-CHO*was trained using ZeroCostDL4Mic (22) notebooks. From all the videos acquired for the study, 105 were chosen for the creation of the ground truth. For each video, a random time-point was selected and all the rounded cells (either before or after division) were manually annotated with a unique label (*i*.*e*., instance segmentation). The data was split into 85 and 20 videos for training and testing respectively. The model was trained on a Tesla T4 GPU for 500 epochs with patches of 525 *×* 512 pixels, a batch size of 20, a grid size of 2 *×* 2 for the first convolutional layer of the model, the Mean Squared Error (MSE) loss function and an initial learning rate of 5 *·* 10^*≠*3^. Data augmentation composed of rotations, translations and mirroring was also applied.Before the training, all the images were downsampled to a pixel size of 0.865 *µu ×* 0.865 *µm* (*i*.*e*., downsampling of a factor of 1.5709), so the size of a cell in the mitotic rounding in pixels is smaller than the input size used to train StarDist. This resampling factor was chosen to ensure that the receptive field of the U-Net in StarDist covers a wide enough region to identify the cell contours characteristic of the rounded cell. The accuracy results in the test set were as follows: intersection over union (IoU) (0, 530), false positive (6.05), true positive (17, 05), false negative (4.6), precision (0.6712), recall (0.790), accuracy (0.555), f1 score (0, 699), true detection (21.65), predicted detections (23.1), mean true score (0.712), mean matched score (0.901), panoptic quality (0.628). The output PDF reports of ZeroCostDL4Mic for the model training and the quality control check are attached to the supplementary material.

##### D.3. Cell activity estimation

Cell activity was calculated as the difference between consecutive pre-processed frames. First, a normalisation of the brightfield microscopy videos was applied to ensure a uniform and comparable illumination intensity across the entire video: (1) the image intensity values of each frame were normalised with the min-max projection to the [0, 1] range, (2) a bleach correction to remove the illumination artefacts was computed by applying a large low-pass filter (a wide Gaussian filter) and subtracting it from the original image, (3) the image intensity values of the entire video are normalised again with the min-max projection to the [0, 1] range.Second, the contrast of the video is enhanced by applying a Contrast-Limited Adaptive-Histogram Equalization (CLAHE) using the Skimage Python package with a kernel of size 25, a clip limit of 0.01 and 256 bins. Third, we applied a Gaussian filter with *‡* =1 to smooth all the frames in the video and alleviate the impact of the noise in the images. Finally, for each time point *t* in the video, the general cell activity was computed as the difference between pairs of temporal frames as follows:

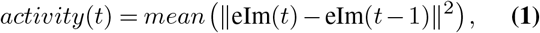

where eIm is a normalised, contrast-enhanced and smoothed image.

#### E. Quantitative data analysis

The percentage of cells in the field of view identified by *StarDist-CHO* at each time point *t* (C(t)) provides a distribution of mitotic rounding events across time. C(t) is calculated by dividing the number of *StarDist-CHO* detections by the number of nuclei detected by *StarDist-versatile*. The peak of cell division is determined as

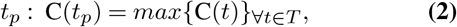

with *t*_*p*_ being estimated for each video, *V*. Cell size was monitored using the instance segmentations from *StarDist-CHO* (Figure 2c-d).

The instance segmentation of *StarDist-CHO* allows for the estimation of the cell size (*S*) (*i.e*., the sum of all the pixels forming the cell mask), which is then used to estimate the cell diameter *D* by resolving the equation

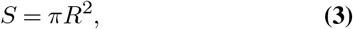

where *R* is the cell radius and *D* = 2*R*. Tracking the distribution of *D* across time we distinguish two populations: mother and daughter cells with a diameter of 20 *µm* and 15 *µm* respectively (Figure 2b,e). This is achieved by classifying all the detections with a diameter larger than 18 *µm* as mother cells and daughter cells otherwise (Figure 2f).

The total *n* number of this analysis is the following: for 385 *nm* is 50 FOVs across 5 replicas for unsynchronised cultures, 50 FOVs across 5 replicas for 0*J/cm*^2^, 20 FOVs across 2 replicas for 0.6*J/cm*^2^, 30 FOVs across 3 replicas for 6*J/cm*^2^ and 30 FOVs across 3 replicas for 60*J/cm*^2^). For 475 *nm*is 37 FOVs across 5 replicas for unsynchronised cultures, 42 FOVs across 5 replicas for 0*J/cm*^2^, 30 FOVs across 3 replicas for 0.6*J/cm*^2^, 27 FOVs across 4 replicas for 6*J/cm*^2^ and 14 FOVs across 2 replicas for 60*J/cm*^2^). For 630 *nm* is 44 FOVs across 5 replicas for unsynchronised cultures, 46 FOVs across 5 replicas for 0*J/cm*^2^, 12 FOVs across 2 replicas for 0.6*J/cm*^2^, 44 FOVs across 5 replicas for 6*J/cm*^2^ and 24 FOVs across 3 replicas for 60*J/cm*^2^). Cell activity at each time point is hardly difficult to compare due to the stochasticity of cell movement at short time windows. Therefore, we calculate the cumulative cell activity after *T* minutes given as

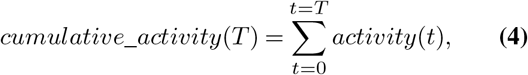

The resulting value at *T* = 420 *min* (7 *h*) is shown in (Figure 2g).

The total *n* number of this analysis is the following: for 385 *nm* is 50 FOVs across 5 replicas for 0*J/cm*^2^, 20 FOVs across 2 replicas for 0.6*J/cm*^2^, 29 FOVs across 3 replicas for 6*J/cm*^2^ and 29 FOVs across 3 replicas for 60*J/cm*^2^). For 475 *nm* is 36 FOVs across 4 replicas for 0*J/cm*^2^, 21 FOVs across 3 replicas for 0.6*J/cm*^2^, 18 FOVs across 2 replicas for 6*J/cm*^2^ and 4 FOVs across 1 replicas for 60*J/cm*^2^). For 630 *nm* is 26 FOVs across 3 replicas for 0*J/cm*^2^, 7 FOVs across 2 replicas for 0.6*J/cm*^2^, 22 FOVs across 3 replicas for 6*J/cm*^2^ and 9 FOVs across 1 replica for 60*J/cm*^2^).

#### F. SYTOX signal quantification

To assess cellular apoptosis, we quantified the percentage of SYTOX-positive cells by dividing the number of cells expressing SYTOX by the total cell count in each field of view. The SYTOX signal was segmented using a two-step process: first, applying Otsu’s thresholding algorithm, followed by a Watershed transformation to delineate individual apoptotic cells. This approach enabled an accurate approximation of apoptotic cell numbers in each frame. The image processing workflow was implemented in Fiji and automated using a custom ImageJ macro for high-throughput analysis. To determine the total cell count, we employed the method described in the previous section, “Estimation of the number of cells in the field of view using virtual cell nuclei staining and segmentation.” A total of 10 FOVs and 1 replica for each condition were assessed.

#### G. Manual annotations of unsynchronised videos

A subset of unsynchronised cells was chosen to identify the delays in mitosis and cell arrest due to photodamage. Those cells entering mitosis were manually labelled by detecting the first time point in which they show the mitotic rounding until the resolvable emergence of two daughter cells. Cells that failed to produce two daughter cells by the end of the timelapse (8 hours) but still attained mitotic rounding are considered to be cell cycle arrested. This annotation lets us distinguish between successful mitoses and cell arrest and calculate the time for cell division or arrest. A total of 3 FOVs per condition were annotated and tracked.

##### Note S1: Setting up and using PhotoFiTT in diverse systems

PhotoFiTT is designed to be a versatile and adaptable framework that can be integrated into a wide range of existing live-cell imaging workflows. Its primary aim is to efficiently evaluate and mitigate phototoxicity. This detailed guide walks researchers through the steps required for implementing PhotoFiTT, reflecting the components illustrated in **Figure 2a**.

#### H. Cell Culture and Synchronisation

The first step in the PhotoFiTT workflow involves preparing the cell samples. This process can be adapted to various cell types, but for optimal results, adherent cell lines capable of division should be used. The following steps can be overviewed in **Figure 2a** diagram blocks: “Unsynchronised cell culture”, “Synchronised cell culture”, “Cell synchronisation drug” and “Drug removal”.

1. **Cell seeding:** Seed cells on an appropriate imaging substrate, such as an 8-well chambered cover glass. The seeding density should be optimised for your specific cell type, but as a starting point, consider using 6 *×* 10^4^ cells per well if using CHO cells. Optimal cell density is crucial for obtaining high-quality images for analysis. Overly dense cell populations can complicate image analysis due to overcrowding and may inhibit cells from re-entering interphase after division due to limited space. Conversely, sparsely populated fields may yield insufficient data for robust statistical analysis.
2. **Synchronisation (optional):** Incubate cells with 10 *nM* RO-3306 (a CDK1 inhibitor) for 16 *≠* 18 hours. This arrests cells at the G2/M boundary, allowing for a coordinated release into mitosis. It is not advisable to incubate cells for longer than 20 hours in RO-3306 as it can induce apoptosis (1).
3. **Unsynchronised option:** After seeding, allow cells to grow for the same duration as the synchronised cell population (16 *≠* 18 hours).

**Recommendation:** Conduct simultaneous experiments with both synchronised and unsynchronised cell populations. This approach allows for the fine-tuning of synchronisation timings and provides insights into the impact of synchronisation on phototoxicity sensitivity for the specific cell type under study.

#### I. Photodamage Irradiation

This phase replicates the illumination conditions encountered by cells in fluorescence microscopy experiments, corresponding to the “Phototoxicity Irradiation” segment depicted in **Figure 2a**. It’s designed to closely mimic the intensity and duration of light exposure typical in such experiments, providing a realistic assessment of potential phototoxic effects.

1. **Microscope Configuration:** Employ a fluorescence microscope outfitted with light sources that can replicate the specific illumination conditions required for assessing phototoxicity. This setup is crucial for accurately simulating the light exposure patterns cells will experience during the experiments.
2. **Power Calibration Protocol:** Initiate each experimental session by calibrating the photodamage irradiation light’s intensity. This step is paramount to guarantee uniformity in experimental conditions. Employ a power meter to accurately measure the light intensity (*W/cm*^2^) at the sample plane, ensuring the microscope’s internal chamber has stabilised at 37°*C*. Control for the fact that light intensity can vary over time. This calibration is critical before commencing each experiment to maintain consistency and reliability in results.
3. **Illumination Conditions:** Configure several fields of view within a single well, assigning unique illumination conditions to each well. This strategy enables the simultaneous evaluation of various light exposure scenarios, facilitating a comprehensive analysis of phototoxic effects across different intensities and durations.
4. **Exposure Process**
  - **Area Selection:** Ensure the selection of distinct, non-overlapping fields for each exposure condition to prevent double irradiation of the same area. Avoid tiling patterns that might lead to such overlaps.
  - **Location Tracking:** Record the coordinates of each illuminated area. This information is crucial for subsequent time-lapse imaging, ensuring accurate follow-up on the exposed regions.
  - **Illumination Application:** Implement the predefined illumination protocols precisely for each designated area, adhering to the established parameters for light intensity and exposure duration.
  - **Media Change for Synchronised Cells:** For cells undergoing synchronisation, wash thoroughly with PBS twice before media replacement. We use a custom-designed 3D-printed multi-syringe adaptor (available on the PhotoFiTT GitHub repository) for efficient and uniform media changes across multiple wells simultaneously. This adaptor is compatible with an 8-well chamber, allowing for simultaneous media changes in 4 wells (**Figure S3**). The consistency of the media exchange process is important for accurate comparison of cell division timings across different experimental setups. Consider that the interval between synchronisation drug removal and the onset of mitotic cell rounding is typically under 10 minutes.

#### J. Live-cell Imaging

Following the completion of the light irradiation stage, the experimental protocol transitions to the timelapse imaging phase. This critical step, denoted as “Acquisition” in the schematic presented in **Figure 2a**, involves capturing sequential images over time to monitor and document the cellular responses to the previously applied light conditions.

1. **Media Exchange Protocol:** Transition the cells to an imaging-optimised medium to enhance both image quality and cell health during observation. Use phenol-red free Fluorobrite DMEM, fortified with 4 *mM* GlutaMAX for sustained cellular metabolism, 42 *µM* gentamicin to prevent microbial contamination, and 10% FBS to support cellular growth and viability.
2. **Microscope Configuration:**
  - Use a high-quality objective that allows to resolve a large number of cells.
  - Ensure the imaging chamber is regulated to maintain an environment that mirrors physiological conditions, with a constant temperature of 37°*C* and a 5% CO_2_ concentration.
3. **Acquisition Parameters:**
  - Use low-intensity brightfield illumination to minimise phototoxicity.
  - Schedule image acquisition over a span of 6 *≠* 8 hours, with images captured every 4 minutes. This frequency balances the need for detailed temporal resolution while constraining phototoxicity.
  - Select a magnification that allows for clear visualisation of individual cells and their mitotic stages (e.g. a 20*X* magnification objective).
4. **Multi-Position Imaging Protocol:** Configure the microscope for automated imaging of all areas subjected to light exposure, as well as designated control regions. For experiments incorporating a tiling strategy, align the regions of photodamage irradiation with those selected for detailed imaging.

**Recomentation:** Implement adaptive focus control, if available, to ensure stable focus over extended imaging sessions. Cellular focus can vary, especially between interphase cells and those in mitotic rounding or division. Identify cells at the mitotic rounding stage to fine-tune the focus, aiming for a balance that accommodates both cell states effectively. This adjustment should be completed prior to initiating photodamage irradiation and drug removal processes, to prevent any delays in live-cell imaging that could impact the integrity of the experimental results.

#### K. Image Analysis Workflow

The PhotoFiTT image analysis pipeline comprises several steps to extract quantitative data from time-lapse images. This workflow integrates deep learning techniques with traditional image processing methods to provide a comprehensive analysis of cellular behaviour under various photodamage irradiation conditions. These steps correspond to the workflow blocks “Data annotation”, “Identification”, “Virtual nuclei staining”, “Cell nuclei segmentation”, “Mitotic rounding detection”, “Calculation of cell activity” illustrated in **Figure 2a**

1. **Cell Detection and Quantification (deep learning-based image analysis):** We used virtual staining. Alternatively, one could use existing pipelines or pre-trained models for cell segmentation when available. This processing is only applied to the first time point of each video.
  - **Virtual Staining:** Apply a Pix2Pix model or an alternative deep learning approach to generate virtual nuclear stains from brightfield images. This step is crucial for label-free cell detection and is performed only on the first frame of each video sequence. See Methods section B for the experimental acquisition of ground truth data and Methods section D for the model training. When reproducing our setup, use our pre-trained Pix2Pix model.
  - **Nuclei Segmentation:** Use the pretrained *StarDist-versatile* model provided by the original authors to segment individual nuclei in the virtually stained images, providing accurate cell counts and positions.
  - **Initial Cell Quantification:** The number of detected nuclei serves as the baseline cell count for each field of view, enabling tracking of population dynamics over time.
2. **Mitotic Cell Identification (deep learning-based image analysis):**
  - **CHO-specific Detection:** For Chinese Hamster Ovary (CHO) cells, employ the specialised StarDist-CHO model to identify cells in the mitotic rounding phase, leveraging its training on the unique morphology of dividing CHO cells.
  - **For other cell lines:** For other cell types or experimental conditions, manually annotate a representative image set and train a new StarDist model following the protocol outlined in Methods Section D.
3. **Cell Size Analysis and Classification:**
  - **Morphometric Measurements:** Exploit the instance segmentations from StarDist-CHO to calculate cell areas and estimate cell diameters.
  - **Cell Stage Classification:** Categorise cells as “mother” (diameter *>* 18 *µm*) or “daughter” (diameter *>* 18 *µm*) based on their size, enabling tracking of cell division progression.
4. **Quantification of Cellular Activity:**
  - **Image Preprocessing:** Enhance brightfield images through illumination normalisation, noise reduction, and contrast enhancement to optimise subsequent analysis.
  - **Dynamic Activity Measurement:** Compute frame-to-frame differences to quantify overall cellular activity, capturing subtle changes in cell morphology and position.
  - **Cumulative Activity Assessment:** Calculate the total cellular activity over the entire imaging period to assess long-term effects of light exposure.

**Implementation Guidelines:**

1. **Deep Learning-based Image Processing:**
  - **Local Processing:** Install DL4MicEverywhere (2) and execute the appropriate notebooks for Pix2Pix and StarDist models.
  - **Cloud-based Processing:** Use ZeroCostDL4Mic (3) notebooks on Google Colab for Pix2Pix and StarDist implementations.
  - **Pre-trained Models:** Access our validated models from Zenodo for immediate use or as starting points for transfer learning.
2. **Data Analysis and Processing:**
  - **Environment Setup:** Configure the required Python environment as specified in the PhotoFiTT repository.
  - **Data Organisation:** Structure raw videos and generated masks according to the provided template.
  - **Analysis Execution:** Employ the provided Jupyter Notebooks to process data and replicate the analysis pipeline.
3. **Quality Control and Validation:**
  - **Manual Verification:** Manually check a subset of images to ensure accurate cell detection and classification.
  - **Reproducibility Assessment:** Compare results across experimental replicates to ensure consistency and reliability.

#### L. Data Interpretation and Optimisation

The final phase of the PhotoFiTT workflow involves interpreting the results and using them to refine your imaging protocols. The following analysis is supported by the output of PhotoFiTT’s Jupyter notebooks. It represents the “Data analysis” workflow illustrated in **Figure 2a**

1. **Mitotic Timing Analysis:**
  - Plot the distribution of mitotic rounding events over time for each condition.
  - Identify the peak division time for control populations (typically around 50 minutes post-synchronisation release).
  - Quantitatively assess delays in this peak for different light exposures and wavelengths.
2. **Cell Size Dynamics Evaluation:**
  - Monitor the temporal evolution of mother and daughter cell proportions.
  - Identify delays in daughter cell emergence, indicative of division slowdown.
  - Identify conditions leading to persistent large cell populations, suggesting potential cell cycle arrest.
3. **Cellular Activity Assessment:**
  - Perform a comparative analysis of cumulative activity levels across different experimental conditions.
  - Interpret reduced activity as a potential indicator of cellular stress or compromised viability.
4. **Phototoxicity Threshold Determination:**
  - Identify the minimum light dose that induces detectable alterations in mitotic timing, cell size distribution, or cellular activity.
  - Use these thresholds as illumination constraints for designing imaging protocols.

##### Note S2: Optimising Live-Cell Imaging Experiments with PhotoFiTT

To maximise the effectiveness of PhotoFiTT in optimising live-cell imaging experiments, follow these comprehensive steps:

1. **Establish a Baseline:**
  - Conduct initial assessments using a wide range of photodamage irradiation conditions relevant to your typical experiments.
  - This baseline will provide crucial insights into where your current protocols fall on the phototoxicity spectrum.
2. **Iterative Protocol Optimisation**
  - Use PhotoFiTT results to guide the fine-tuning of acquisition parameters.
  - Implement multiple rounds of optimisation as needed, systematically testing new settings.
3. **Continuous Monitoring:**
  - Integrate PhotoFiTT assessments into routine workflow, particularly when:
  - Introducing new cell types or modifying culture conditions.
  - Incorporating non-standard drugs or chemicals into your experiments.
  - Adopting novel fluorophores or labelling strategies.
  - Implementing new imaging modalities or upgrading hardware.
4. **Informed Experimental Design:**
  - Leverage PhotoFiTT data to enhance experimental planning:
    - Determine the maximum allowable number of time points or z-stacks before phototoxicity becomes a significant factor.
    - Assess the feasibility of photoactivation or optogenetic experiments based on cellular light sensitivity.
    - Optimize the sequence of multi-colour imaging to minimize potential damage from shorter wavelengths.
5. **Reporting and Reproducibility:**
  - Include detailed PhotoFiTT-derived phototoxicity assessments in your methods sections and supplementary data. This quantitative approach to evaluating phototoxicity will improve the reproducibility and reliability of your imaging studies.

---

**Fig. S1.**
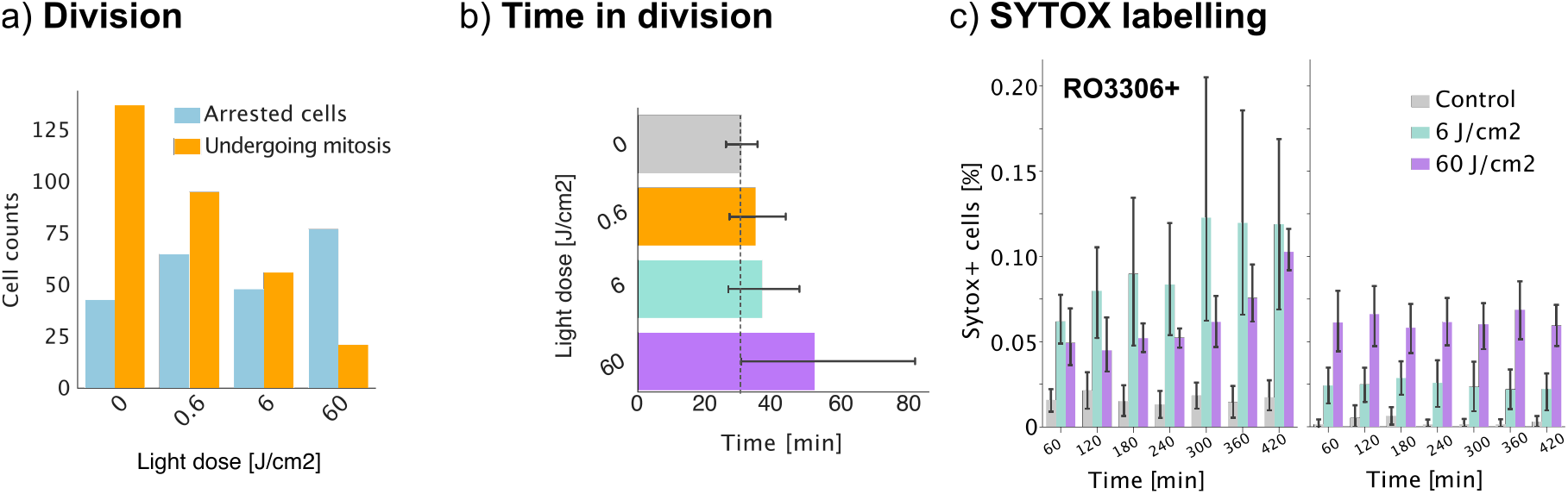
Impact of phototoxicity on non-synchronised cell populations. Figure presents the effects of phototoxicity on non-synchronised adherent mammalian (Chinese Hamster Ovary CHO) cell populations under near-UV (385 *nm*) light exposure. a) Relationship between light dose and cell fate, showing a decrease in cell division and an increase in cell arrest with higher UV doses. b) Dose-dependent delay in cell division is quantified by measuring the time from mitotic rounding to the emergence of two distinct daughter cells, revealing longer delays at higher near-UV doses. c) Comparison of apoptotic cell rates (indicated by SYTOX-positive staining) between synchronised and non-synchronised cell populations under near-UV exposure highlights higher vulnerability of synchronized populations to phototoxic damage.

**Fig. S2.**
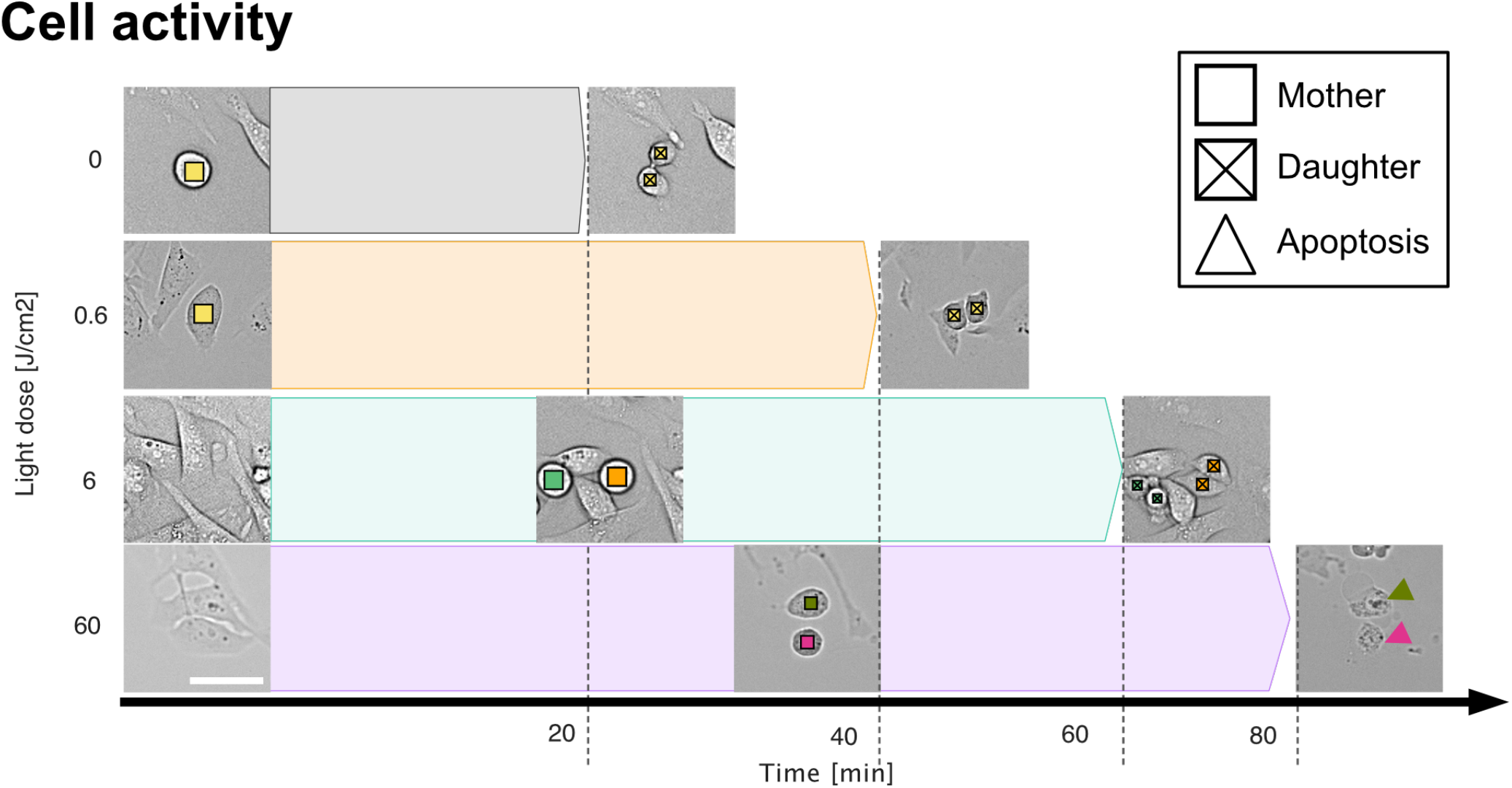
Cell activity after light exposure. Temporal illustration of near-UV (385 *nm*) light dose effects on cell division timing, arrest, and apoptosis. In the absence or under low light exposure, mother cells (square) proceed to divide resulting in the formation of daughter cells (square with a cross inside). Following high light exposure, cells present delays in mitotic events resulting in delays or even apoptosis (triangle). Scale bar 50 *µm*.

**Fig. S3.**
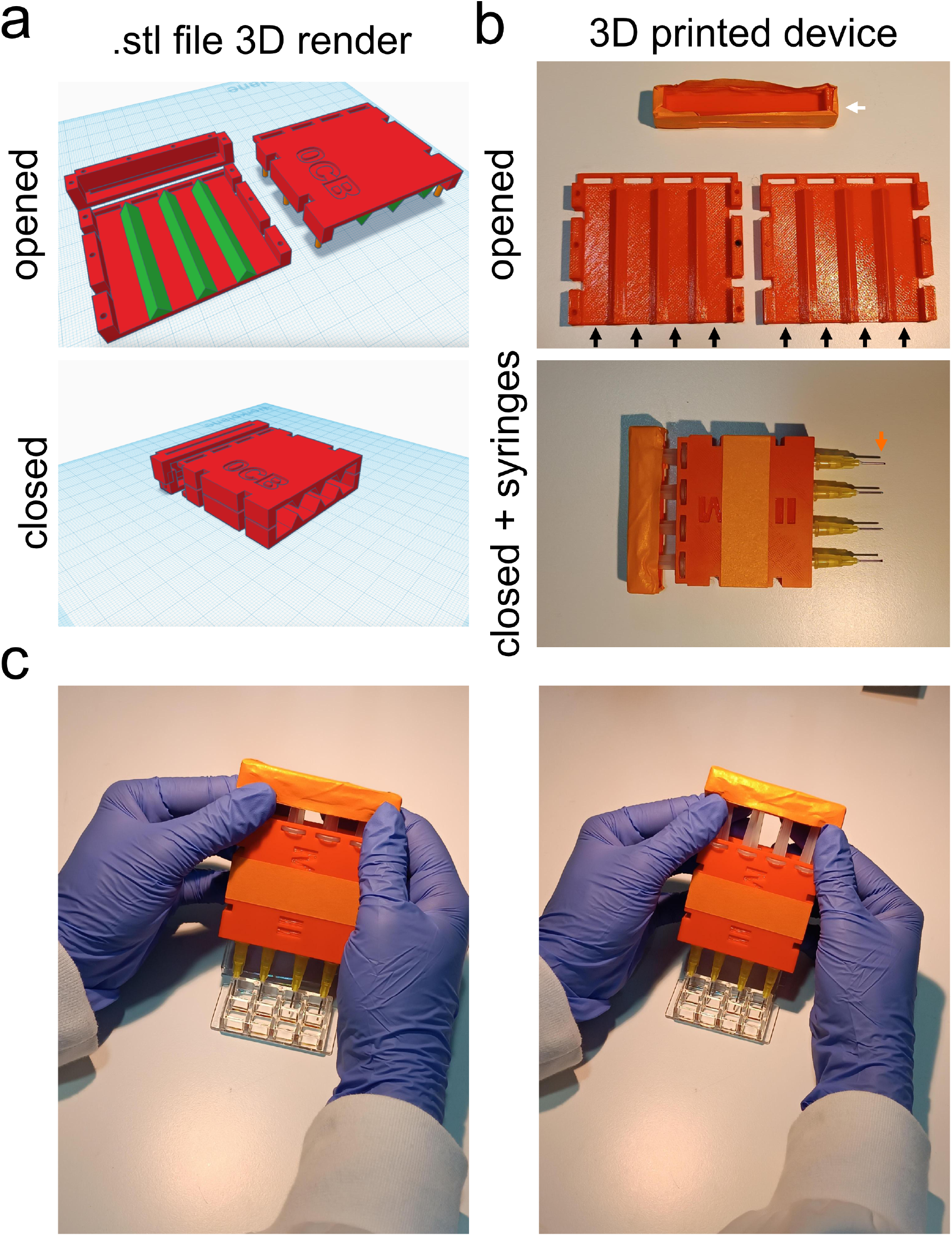
3D-printed multi-well syringe adaptor for efficient media changes. This custom device accommodates four 2.5 mL syringes for use with standard 8-well glass-bottom imaging chambers. a) 3D render of the .stl file in open and closed configurations. b) 3D-printed device with and without syringes. Black arrows indicate well spacing matching the 8-well chamber; white arrow shows the plunger thumb rest for uniform syringe motion; orange arrow points to recommended blunt needles for optimal media uptake. c) Demonstration of the adaptor’s use on an 8-well chamber. The device was designed using TinkerCAD and printed with a Prusa MK4 3D printer. Additional securing with tape is optional.

**Table S1.** Irradiation ranges and microscopy modalities for different fluorescence microscopy techniques. This table summarises the typical irradiation ranges and average irradiation intensities for various fluorescence microscopy techniques, along with corresponding references. STED: Stimulated Emission Depletion; SMLM: Single Molecule Localization Microscopy; RESOLFT: Reversible Saturable Optical Fluorescence Transitions; SRRF: Super-Resolution Radial Fluctuations; TIRF: Total Internal Reflection Fluorescence; SIM: Structured Illumination Microscopy; LLS: Lattice Light-Sheet.

| **Irradiation range**<br>(W/cm<sup>2</sup>) | **Irradiation average**<br>(W/cm<sup>2</sup>) | **Microscopy modality** | **References** |
| --- | --- | --- | --- |
| 1000 - 20000 | 10000 | STED | Wildanger et al., 2008 |
| 1000 - 10000 | 5000 | SMLM/RESOLFT | Chen et al., 2018;<br>Grotjohann et al., 2011 |
| 100 - 5000 | 1000 | Confocal | Icha et al., 2017 |
| 50 - 1000 | 100 | SRRF | Culley et al., 2018 |
| 5 - 100 | 10 | TIRF; SIM | Kwakwa et al., 2016;<br>Li et al., 2015 |
| 0.5 - 100 | 5 | LLS; Wide-field | Icha et al., 2017;<br>Schermelleh et al., 2019 |

**Video S1: Detailed procedure for the PhotoFiTT experiment setup and image acquisition**. This instructional video outlines the PhotoFiTT experimental setup and data acquisition process. Initially, standard adherent cells are seeded in suitable culture dishes. These cells are then synchronised using the CDK1 inhibitor, RO-3306, for a period of 16 to 18 hours to ensure uniform cell cycle progression. Prior to imaging, the microscope’s excitation light intensity is calibrated to maintain consistency across all experiments. The cells are exposed to a predetermined dose of excitation light, simulating conditions encountered during fluorescence microscopy. Following light exposure, the cells are washed with PBS to remove any residual inhibitor, preparing them for live imaging. The video concludes with the initiation of live-cell acquisition, capturing the dynamic responses of cells to phototoxic stress.

**Video S2. Demonstrating the PhotoFiTT analytical workflow**. This video provides a comprehensive walkthrough of the PhotoFiTT analytical process, utilising the Jupyter notebooks available in the PhotoFiTT GitHub repository (https://github.com/HenriquesLab/PhotoFiTT). The demonstration covers each step in detail, showcasing how to effectively use the provided tools for analysing phototoxicity effects on cells.

**Video S3. Dynamics of adherent cells after light exposure**. Synchronised CHO cells following 385 nm (near-UV) light exposure. Non-exposed cells present a clearly defined peak in mitotic rounding and cell division (white arrow). Cells exposed to a dose of 0, 6*J/cm*^2^ present a slight delay but can complete division (orange arrow). Cells exposed to a dose of 6*J/cm*^2^ present a delay in mitotic rounding. In most cases, the cells present aberrant division (blue arrow) or are unable to complete division (teal arrow). Cells exposed to the high dose of 60*J/cm*^2^ become arrested and can result in cell death (purple arrows).

